# Integrative transcriptomic and metabolic analyses of the mammalian hibernating brain identifies a key role for succinate dehydrogenase in ischemic tolerance

**DOI:** 10.1101/2023.03.29.534718

**Authors:** Joshua D. Bernstock, Cory M. Willis, Monica Emili Garcia-Segura, Edoardo Gaude, Daniela Anni, Yan-ja Lee, Luke W. Thomas, Alva Casey, Nunzio Vicario, Tommaso Leonardi, Alexandra M. Nicaise, Florian A. Gessler, Saef Izzy, Mario R. Buffelli, Jakob Seidlitz, Shriya Srinivasan, Michael P. Murphy, Margaret Ashcroft, Marco Cambiaghi, John M. Hallenbeck, Luca Peruzzotti-Jametti

## Abstract

Ischemic stroke results in a loss of tissue homeostasis and integrity, the underlying pathobiology of which stems primarily from the depletion of cellular energy stores and perturbation of available metabolites^1^. Hibernation in thirteen-lined ground squirrels (TLGS), *Ictidomys tridecemlineatus*, provides a natural model of ischemic tolerance as these mammals undergo prolonged periods of critically low cerebral blood flow without evidence of central nervous system (CNS) damage^2^. Studying the complex interplay of genes and metabolites that unfolds during hibernation may provide novel insights into key regulators of cellular homeostasis during brain ischemia. Herein, we interrogated the molecular profiles of TLGS brains at different time points within the hibernation cycle via RNA sequencing coupled with untargeted metabolomics. We demonstrate that hibernation in TLGS leads to major changes in the expression of genes involved in oxidative phosphorylation and this is correlated with an accumulation of the tricarboxylic acid (TCA) cycle intermediates citrate, cis-aconitate, and α-ketoglutarate-αKG. Integration of the gene expression and metabolomics datasets led to the identification of succinate dehydrogenase (SDH) as the critical enzyme during hibernation, uncovering a break in the TCA cycle at that level. Accordingly, the SDH inhibitor dimethyl malonate (DMM) was able to rescue the effects of hypoxia on human neuronal cells *in vitro* and in mice subjected to permanent ischemic stroke *in vivo*. Our findings indicate that studying the regulation of the controlled metabolic depression that occurs in hibernating mammals may lead to novel therapeutic approaches capable of increasing ischemic tolerance in the CNS.

---

Stroke is one of the largest contributors to morbidity/mortality worldwide and continues to increase in both incidence and prevalence^3^. From a pathophysiological perspective, ischemic stroke sets in motion a complicated array of metabolic and neurophysiological perturbations, which ultimately lead to permanent CNS damage^4^. Because of the inherent complexity of this pathological process, a sensible approach to treatment must centre on the targeting of regulators of CNS homeostasis in an effort to improve outcomes after permanent brain ischemia. In this sense, model organisms that have evolved homeostatic and cytoprotective processes to promote a natural tolerance to ischemic conditions may prove invaluable.

TLGS are one of several types of mammals with the physiological capability of undergoing hibernation in response to extreme environmental conditions^5^. TLGS hibernation cycle is a tightly regulated process that can be studied by following the molecular modifications that occur as they progress through several phases (see **Methods** for detailed overview of the TLGS hibernation cycle). This process begins with a physiological baseline state defined as active cold room (ACR) and progresses to the entrance of hibernation (ENT) phase. From there, the TLGS proceed to early hibernation (E_Hib) and finally the late hibernation/torpor (L_Hib) phase. During L_Hib, cerebral blood flow is severely diminished (i.e., a ~90% reduction from baseline) and reaches values comparable to those seen in the ischemic core of stroke patients^2,6^. Nonetheless, no ischemic damage nor functional deficits are described in these animals after prolonged periods of hypoxia spent in L_Hib^2,7^. Notably, the neuroprotective phenotype associated with hibernation not only allows neural tissue to cope with extreme conditions, but also appears to promote increased resistance to superimposed neural injury^8^.

The goal of this study was to comprehensively interrogate the combined gene expression and metabolic profiles of TLGS brain tissue at different stages of hibernation via RNA sequencing integrated with ultrahigh performance liquid chromatography-tandem mass spectroscopy (UPLC-MS/MS)-based metabolomics in an effort to identify new targets capable of promoting tolerance to brain ischemia (**Fig. 1a**).

**Figure 1.**
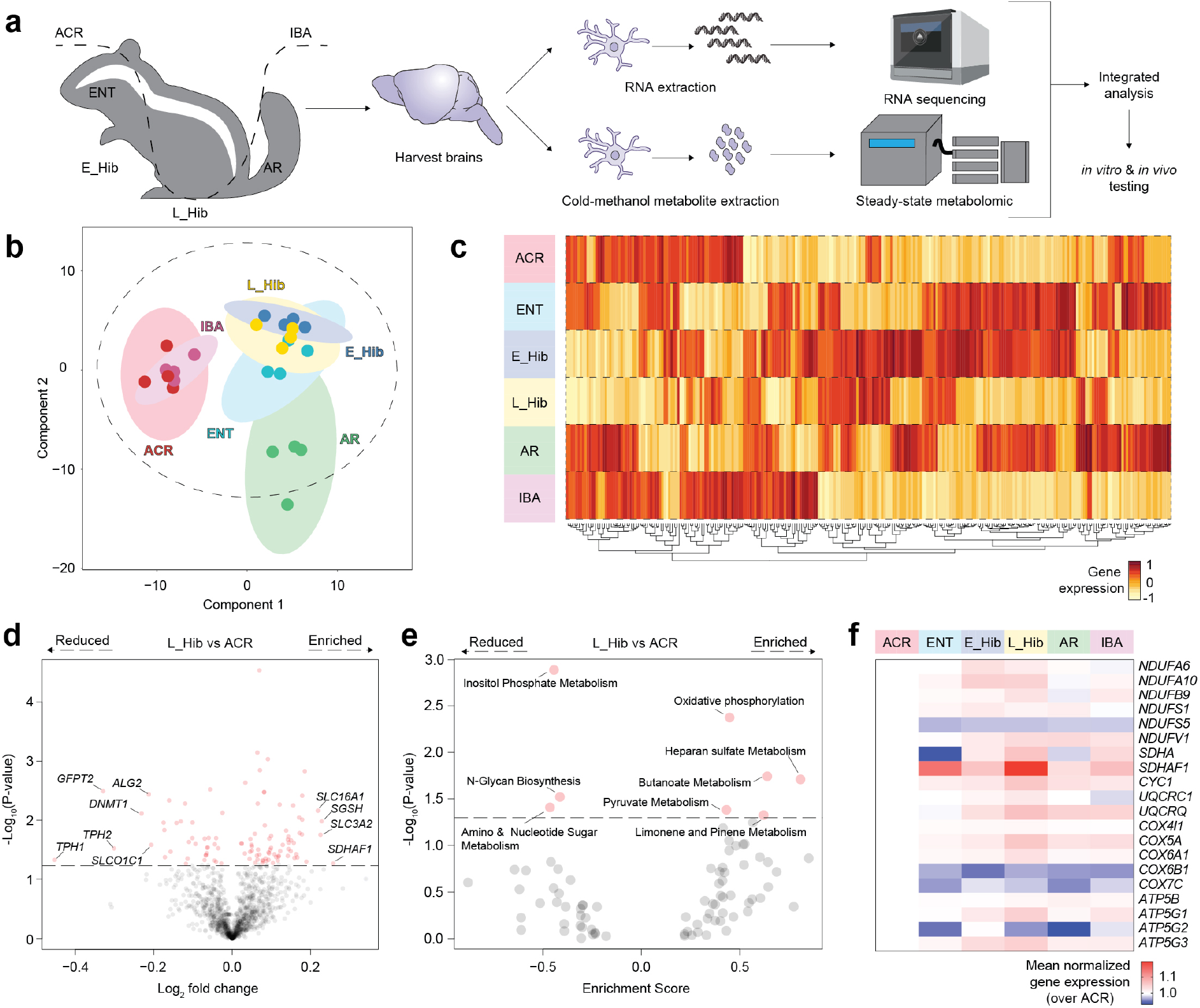
Gene expression and pathway enrichment analyses. **a,** Schematic overview of the workflow for steady-state global RNA sequencing and metabolomics profiling of brains obtained from thirteen-lined ground squirrels (TLGS) at defined stages of hibernation. **b**, Sparse Partial Least Squares - Discriminant Analysis (sPLS-DA) of metabolic genes at defined stages of hibernation. Individual points indicate each biological replicate, n=4 biological replicates per group. **c**, Heat map representation of the pathways (Wiki Pathways) found significantly enriched by GSEA analysis in each stage of hibernation. **d**, Volcano plot showing the pairwise comparison of the metabolic genes between L_Hib and ACR, n=4 biological replicates per group. **e,** Volcano plot showing the significantly enriched metabolic pathways according to GSEA analysis performed between L_Hib and ACR. **f**, Heat map of metabolic genes (P-value < 0.1) defining the oxidative phosphorylation pathway at defined stages of hibernation. Abbreviations: ACR (active cold room), ENT (entrance of hibernation), E_Hib (early hibernation), L_Hib (late hibernation/torpor), AR (arousal), IBA (interbout arousal).

Sparse partial least squares - discriminant analysis (sPLS-DA) performed on the n=18,492 genes identified via RNA sequencing showed a clear separation between hibernating TLGS (i.e. ENT, E_Hib, L_Hib, and arousal-AR) *vs* control animals (i.e., ACR and interbout arousal-IBA) (**Fig. 1b**). Based on gene set enrichment analysis (GSEA), we found that ~17% of the significantly enriched pathways belonged to metabolic functions, highlighting the involvement of cellular metabolism in the mammalian hibernation cycle (**Fig. 1c, Table 1**). We also found that each stage of hibernation was defined by predominant alterations of specific gene pathways, with mitochondrial complex II (succinate dehydrogenase, SDH) assembly being the most relevant GSEA pathway in L_Hib (**Table 1**).

We next focused on the subset of genes with known metabolic functions (n=1,517) to identify regulators of the controlled metabolic depression that occurs during L_Hib (*vs* control ACR) and protects against hypoxic damage. We found *SDHAF1, SLC3A2, SGSH* and *SLC16A1* to have the highest upregulation among the differentially expressed genes (DEGs) in L_Hib *vs* ACR; while *TPH1*, *GFPT2*, *TPH2*, *DNMT1*, *ALG2* and *SLCO1C1* were the most downregulated DEGs in L_Hib *vs* ACR (**Fig. 1d**, **Table 1**). GSEA analysis of these DEGs showed that *oxidative phosphorylation* was the most significantly upregulated metabolic pathway in L_Hib *vs* ACR, followed by *butanoate, heparan sulfate, pyruvate*, and *limonene and pinene metabolism* (**Fig. 1e**, **Table 1**). Whereas processes involved in *amino and nucleotide sugar metabolism, N-glycan biosynthesis*, and *inositol phosphate metabolism* were significantly downregulated in L_Hib *vs* ACR (**Fig. 1e**, **Table 1**). When we analysed the DEGs determining each metabolic pathway at the different stages of TLGS hibernation (**Fig. 1f**, **Extended data Fig. 1**), we found that *SDHAF1* and *SDHA* were among the most upregulated *oxidative phosphorylation* genes, in particular during L_Hib (**Table 1**). Interestingly, *SDHAF1* encodes for one of the chaperone proteins involved in the assembly of the SDH complex and plays a key role in its protection against oxidative stress^9^. Overall, these data suggest that major alterations to metabolic genes occur during TLGS hibernation, and that oxidative phosphorylation, and in particular SDH-related genes, are at the forefront of these adaptive changes.

To further investigate the involvement of metabolism in the response to hibernation in TLGS, we next analysed the n=462 metabolites identified via UPLC-MS/MS analysis. SPLS-DA showed a clear separation between the individual stages of TLGS hibernation cycle (**Fig. 2a**). This was coupled with clustering of metabolites belonging to several metabolic pathways (**Fig. 2b**, **Table 2**). We next performed pathway enrichment analysis of all the metabolites with VIP score ≥1 (n=185) followed by heuristic multiple linkage clustering of enriched pathways to provide a comprehensive view of the metabolic changes that occur during the TLGS hibernation cycle (**Table 2**). We excluded pathways that (i) did not cluster or (ii) were annotated into multiple clusters, thus identifying a total of n=12 unique enrichment clusters (**Fig. 2c**), which included key biological processes (**Extended data Fig. 2, Table 2**). The most biologically relevant cluster (based on statistical significance and elevated median metabolite ratio^10^) was *enrichment cluster 6*, which included metabolites involved in *alanine and aspartate metabolism*, the *TCA cycle*, and *amino acid metabolism* (**Fig. 2d, Table 2**).

**Figure 2.**
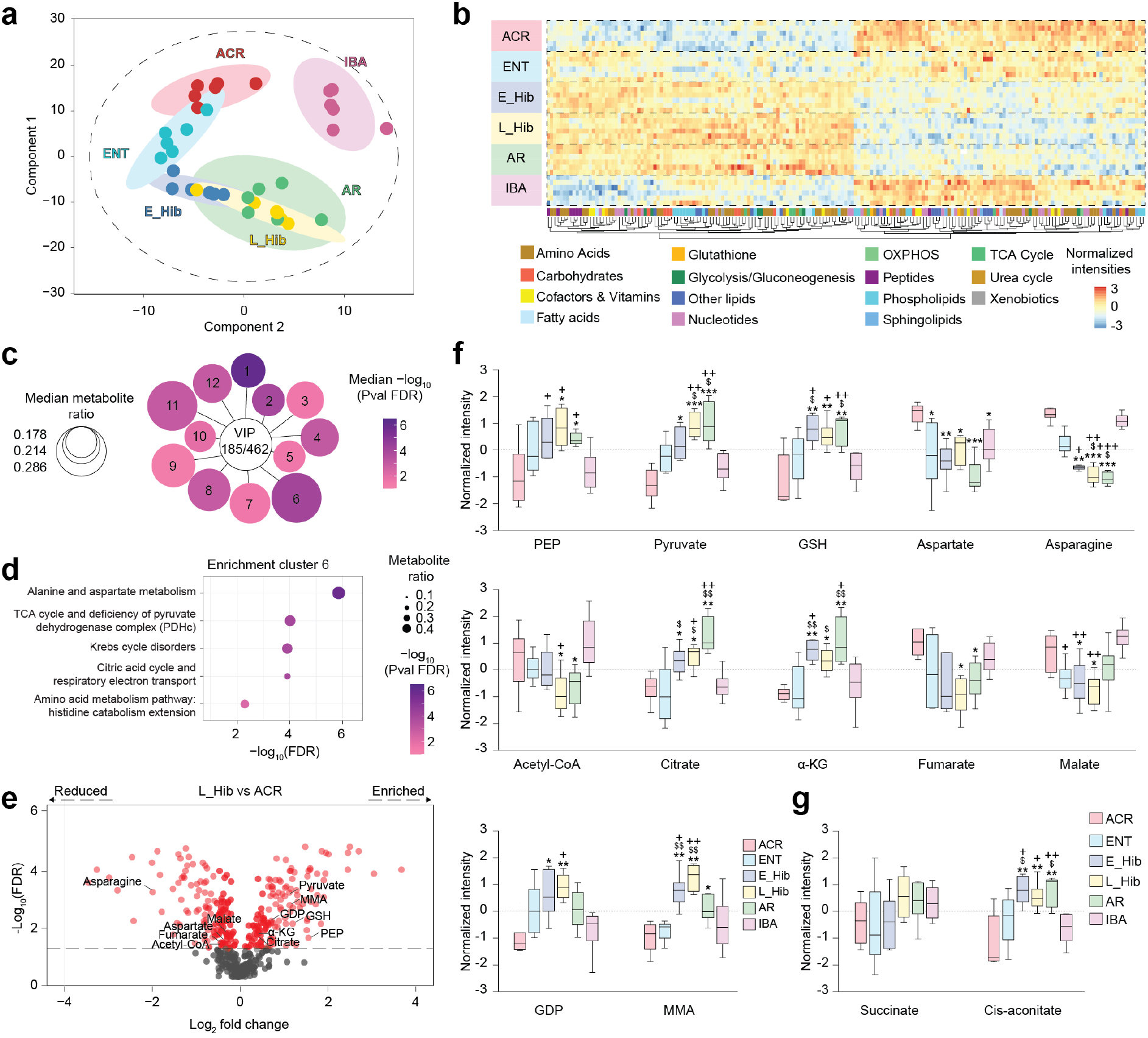
Metabolic profiling of hibernators show key TCA cycle alterations. **a**, Sparse Partial Least Squares - Discriminant Analysis (sPLS-DA) of the global metabolomics data obtained via UPLC-MS/MS of the hibernators’ brains at defined stages of hibernation. Individual points represent each biological replicate, n=6 biological replicates per group. **b**, Heat map of normalized intensities showing clustering of the 462 metabolites identified in the hibernators’ brains, n=6 biological replicates per group. **c**, Diagram showing the clusters derived from pathway enrichment analysis of the 185/462 metabolites with VIP score ≥1. Circle size is proportional to the median metabolite ratio in the pathway and colorimetric scale shows the median −log10 value of Benjamini-Hochberg (BH) corrected significance, n=6 biological replicates per group. **d,** Individual enrichment analysis of biological pathways assigned to *enrichment cluster 6*. Circle size is proportional to the metabolite ratio in the pathway and colorimetric scale shows the −log10 value of BH adjusted significance. **e**, Volcano plot showing the pairwise comparison between L_Hib and ACR. Metabolites belonging to *enrichment cluster 6* are shown, n=6 biological replicates per group. **f**, Normalized intensities of the single metabolites belonging to *enrichment cluster 6* at defined stages of hibernation. Significance was determined via Kruskal-Wallis non-parametric testing followed by BH correction for multiple testing. *: p<0.05; **: p<0.01; ***: p<0.001 vs ACR. $: p<0.05; $$: p<0.01 vs ENT. +: p<0.05; ++: p<0.01; +++: p<0.001 vs IBA, n=6 biological replicates per group. **g,** Normalized intensities of succinate and cis-aconitate at defined stages of hibernation, with significance determined via Kruskal-Wallis non-parametric testing followed by BH correction for multiple testing. **: p<0.01 vs ACR. $: p<0.05 vs ENT. +: p<0.05; ++: p<0.01 vs IBA, n=6 biological replicates per group. Abbreviations: ACR (active cold room), ENT (entrance of hibernation), E_Hib (early hibernation), L_Hib (late hibernation/torpor), AR (arousal), IBA (interbout arousal), PEP (phosphoenolpyruvate), GSH (reduced gluthatione), α-KG (α-ketoglutarate), GDP (guanosine-5’-diphosphate), MMA (methylmalonate), SDH (succinate dehydrogenase).

We next focused on the single metabolites constituting *enrichment cluster 6* and their putative role in the controlled metabolic depression that occurs during L_Hib *vs* ACR (**Fig. 2e, Table 2**). *Enrichment cluster 6 m*etabolites linked to glycolytic processes (i.e., phosphoenolpyruvate-PEP, and pyruvate) and glutathione (GSH) were all significantly increased in L_Hib *vs* ACR; while aspartate, asparagine, and acetyl-CoA were significantly decreased in L_Hib *vs* ACR (**Fig. 2f, Table 2**). When focusing on *enrichment cluster 6* metabolites involved in the TCA cycle, we found a dichotomous response to hibernation with a significant increase in metabolites upstream of SDH (i.e., citrate and α-ketoglutarate-αKG) and a decrease of downstream metabolites (i.e., fumarate and malate) in L_Hib *vs* ACR. The SDH related energy metabolite guanosine-5’-diphosphate (GDP) and the dicarboxylic acid methylmalonate (MMA), which has been reported to inhibit SDH activity^11,12^, were also found to be among the *enrichment cluster 6* metabolites that were increased during L_Hib *vs* ACR (**Fig. 2f**). While significant alterations of succinate levels were not captured, we instead found that cis-aconitate (another metabolite known to inhibit SDH activity, but not included in *enrichment cluster 6*)^13^ was also increased in L_Hib *vs* ACR (**Fig. 2g**). Collectively, these data suggest that hibernation in TLGS induces a dichotomy of TCA cycle metabolites that may be explained via an enzymatic block at the level of SDH^14^.

To further confirm the enzymatic targets underlying the controlled metabolic depression that occurs during TLGS hibernation, we integrated the gene expression and metabolomics datasets by identifying the enzyme commission (EC) numbers associated with each significant gene and metabolite. This systems biology approach led to the identification of 330 unique enzymatic functions, out of which only 7 were shared between genes and metabolites (**Fig. 3a, Table 2**). Among these, SDH was identified, which supports our transcriptomic and metabolomic analyses. Altogether these data show that hibernation in TLGS induces major changes in the expression of both genes and metabolites that are linked with key enzymatic functions in the brains of TLGS that increase their tolerance to brain ischemia (**Fig. 3b**).

**Figure 3.**
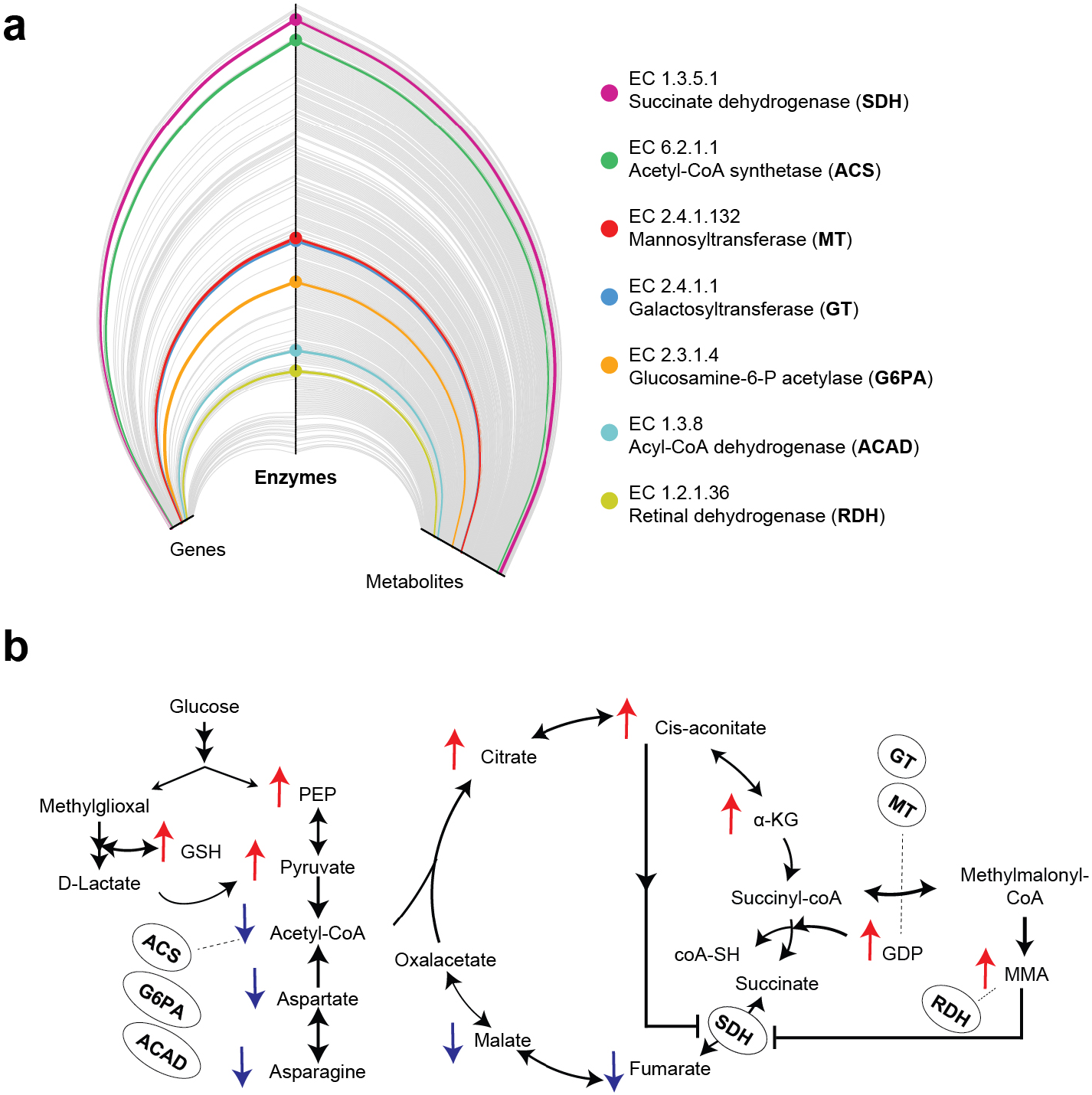
Integrative transcriptomic and metabolic analysis. **a**, Hive plot showing significant (L_Hib *vs* ACR) genes (left axis) and metabolites of *enrichment cluster 6* (right axis) and their associations with enzymatic functions (central axis). Enzymes associated with both significant genes and metabolites are highlighted. **b**, Schematic representation of enzymes (in bold) mapped on the metabolic changes seen in hibernation (directionality is given with red and blue arrows to show *enrichment cluster 6* metabolites that changes in L_Hib *vs* ACR). Abbreviations: EC (enzyme commission).

We next investigated whether SDH inhibition could be induced in relevant cellular and animal models to promote ischemic tolerance, as seen in the hibernating TLGS brain. We first tested the ability of DMM, a molecule capable of crossing cell membranes and the blood brain barrier^15^, to inhibit SDH activity in human SH-SY5Y differentiated neurons (**Fig. 4a**). To mimic ischemic stress *in vitro*, we subjected SH-SY5Y differentiated neurons to hypoxia (1% O2), which was quantified via western blot (i.e., blotting for known hypoxia-induced proteins, such as HIF-1α/HIF-2α, GLUT1, and BNIP3), flow cytometry (for apoptosis/necroptosis), and gene expression analysis (**Extended data Fig. 3**). Upon hypoxia, SH-SY5Y differentiated neurons were treated with DMM (10 mM) or PBS (sham) (**Fig. 4b**). After 24 h we observed a significant shortening of neurites (total length and average length) in SH-SY5Y neurons under hypoxic conditions, which was prevented by DMM treatment (**Fig. 4c-d**).

**Figure 4.**
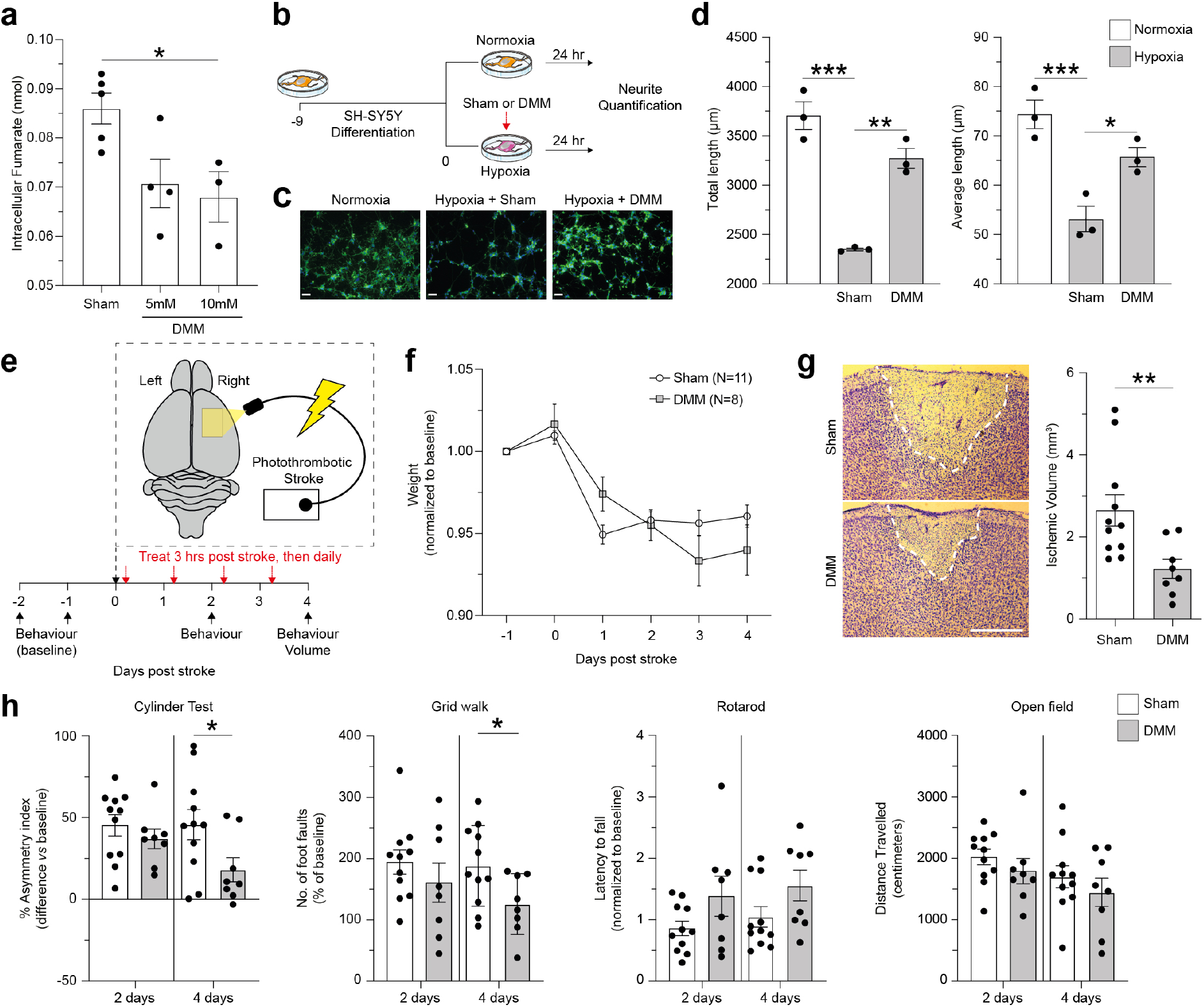
SDH inhibition promotes tolerance to hypoxia/ischemia *in vitro* and *in vivo*. **a,** Treatment with DMM for 1 hr inhibits SDH activity and reduces intracellular fumarate levels in human SH-SY5Y neurons, n=3-5 technical replicates per group. **b**, Schematic overview of the workflow for the *in vitro* experiments on hypoxic SH-SY5Y neurons receiving DMM or sham (PBS) treatment. **c**, Representative pictures of SH-SY5Y neurites under normoxia, hypoxia + sham treatment, and hypoxia + DMM treatment. Green: neurites stained with FITC-phalloidin. Blue: DAPI. Scale bar: 46 μm. **d**, Quantification of total and average neurite length of SH-SY5Y neurons under hypoxia (1% O_2_) after sham/DMM treatment for 24 h. n=3 technical replicates per group. Significance was determined via an ordinary one-way ANOVA followed by Tukey’s multiple comparisons test to correct for multiple comparisons. *: p<0.05; **: p<0.01, ***: p<0.001. **e**, Schematic overview of *in vivo* experiments in the mouse model of photothrombotic stroke receiving DMM or sham (saline) treatment. **f**, Body weight variation after photothrombotic stroke in mice, n≥8 biological replicates per group. **g**, Representative Nissl-stained brain sections showing the lesion in the M1 area and quantification of the ischemic lesion in sham/DMM treated mice. Red dotted lines shows edge of the ischemic lesion, n≥8 biological replicates per group. **: p<0.01 (unpaired t-test). **h**, Behavioural tests specific for the affected forelimb showing a significant amelioration in DMM treated vs sham treated mice with respect to pre-ischemic condition (baseline). DMM treatment resulted in a significant reduction in the asymmetry index (cylinder test) and in the number of foot faults (grid-walk test) with respect to untreated mice, n≥8 biological replicates per group. *: p<0.05 (unpaired t-test). Abbreviations: DMM, dimethyl malonate.

We next employed a model of photothrombotic stroke in the primary motor cortex of mice to test the effects of SDH inhibition on the clinicopathological outcomes of permanent brain ischemia (**Fig. 4e**). This model leads to an occlusion of small cerebral vessels, which causes ischemic cell death and prolonged sensorimotor impairment^16,17^. At 3 h following stroke induction, mice received daily intraperitoneal (IP) injections of either DMM (160mg/kg) or saline (sham) for 4 days. We observed a physiological reduction of body weight at 24 h after stroke induction, which later stabilized and was not statistically significant between groups (**Fig. 4f**). Instead, we found that DMM treated ischemic mice had a significantly lower ischemic lesion volume at 4 days after stroke *vs* sham (**Fig. 4g**). This effect was coupled with a significant recovery of forelimb impairment *vs* sham, as assessed with the cylinder and grid-walk tests (**Fig. 4h**). In a motor coordination test (i.e. rotarod) and in an open field test, both estimating bilaterally symmetric dysfunction, we found a trend toward better outcomes in DMM treated ischemic mice *vs* sham, although this was not significant (**Fig. 4g**). These results confirmed that, in addition to promoting neuroprotection *in vitro*, SDH inhibition promotes ischemic tolerance and prevents functional impairment to permanent brain ischemia *in vivo*.

Previous approaches centred on the detection of metabolic alterations during TLGS hibernation have demonstrated a switch from carbohydrate to lipid metabolism, decreased protein synthesis and protein degradation, and an IBA/AR period-specific increase in gluconeogenesis at the systemic level^18–21^. Tissue specific (i.e., liver) metabolomic analyses of the TLGS have demonstrated seasonal and hibernation cycle variations in purine/pyrimidine metabolism, amino acid, nitrogen, phospholipid catabolism, and lipid metabolism^22,23^. Finally, two recent studies have also shed light on nitrogen recycling during prolonged fasting in the TLGS.^24,25^ In the CNS, a previous study performed a targeted analysis of 18 metabolites via NMR spectroscopy and found changes in neurotransmitters (i.e., GABA increase coupled with glutamate decrease) during torpor^26^. These and other data^27,28^ have laid a foundation for further studies aiming to explore the molecular underpinnings of ischemic tolerance within the hibernating CNS. Yet, no previous body of work has explored the entire cycle and/or have sought to validate the function of ascribed targets in pertinent preclinical models of brain ischemia. Therefore, more comprehensive transcriptomic and metabolomic analyses of the brain from multiple time points during the hibernation cycle was needed to provide novel/actionable insights into the extreme neuroprotection, cytoprotection, and maintenance of homeostasis of which these animals are capable.

Our data constitutes the first comprehensive untargeted survey of the gene and metabolic changes that appear in the hibernating TLGS brain throughout the hibernation cycle. This unique approach allowed us to draw several novel conclusions based on our analyses of hibernation in the TLGS, which, among hibernating mammalians, are one of the most resistant to ischemia. Furthermore, while reperfusion after ischemic stroke is known to increase the risk of oxidative stress and damage^29^, hibernating TLGS are able to exit from hibernation and reinitiate normal metabolic function without incurring CNS damage^30^. This suggests an endowed natural protection against ischemia-reperfusion injury, which could be further explored in the future using our unique dataset. While many transcriptomic and metabolic alterations were differentially regulated during hibernation, we focused on the L_Hib stage of hibernation as this most closely models the conditions seen in the ischemic core of patients. We identified a prominent rearrangement of the TCA cycle, centred at the level of SDH, which appeared to be associated with the natural tolerance to ischemia of hibernating TLGS.

Recent data has shown that succinate accumulates in both the human and mouse brain upon ischemia, and that inhibiting its oxidation via delivery of SDH inhibitors before stroke induction^31^ or before stroke reperfusion^32^ can lead to a dose-dependent decrease in brain injury. Our data build on this body of evidence to unequivocally show that the delayed (i.e., 3 h after stroke) therapeutic systemic administration of DMM is able to significantly ameliorate the clinicopathological outcomes of stroke in the absence of reperfusion. Moreover, we show that DMM has a direct neuroprotective effect on human neurons, where it protects neurites from hypoxic damage. Such effects may ultimately prove transformative in clinical stroke care.

Altogether our data provide a solid basis for future studies aimed at interrogating hibernation, and its related compensatory mechanisms, on a deeper cellular level. This may be key to developing next generation therapeutics for CNS diseases/disorders, including ischemic stroke. By understanding the mechanisms by which hibernators preserve homeostasis during extreme conditions, our work may ultimately be leveraged translationally to influence the care of stroke patients that have suffered permanent ischemic injury.

## Methods

### Hibernation in TLGS

TLGS were captured by USDA-licensed trappers (TLS Research, Bartlett, IL, USA). Experiments were approved by the NINDS Animal Care and Use Committee. Both male and female ground squirrels were used equally for this study and all animals were between one and three years of age (as these animals were caught in the wild, only an age estimate was available). TLGS were housed, fed, and induced to hibernate as described previously^33,34^. Animals in 6 different phases of the hibernation bout (cycle) were differentiated by body temperature (Tb), time, and respiratory rate as previously described^33,34^. ACR: active for 4 to 5 days in the cold chamber with a temperature of 4°C to 5°C (Tb=34°C to 37°C); ENT: entrance of hibernation (Tb=31°C to 12°C); E-Hib: early hibernation (1 day, Tb=5°C to 8°C); L-Hib: late hibernation/torpor (>5 days, Tb=5°C to 8°C); AR: arousal from torpor spontaneously with a respiratory rate >60/min and a persistent low body temperature (Tb=9°C to 12°C); IBA: interbout arousal from a torpor state and returned to a normal metabolic state for at least 7 h (Tb=34°C to 37°C). At each time point, brains were removed, snap-frozen in 2-methylbutane (−50°C), and stored at −80°C until use; n=4 and n=6 animals for each time point were used for transcriptomics and metabolomics studies, respectively (see **Supplementary Information**).

### Photothrombotic stroke experimental design and DMM administration *in vivo*

Male C57/BL6 mice (8–10 weeks of age) were used for this study. All animals were allowed free access to food/water and were maintained in a humidity and temperature-controlled room with a 12-hour light/dark cycle. Procedures were performed according with the EU Council Directive 2010/63/EU on the protection of animals used for scientific purposes. Mice were randomly sub-divided into two experimental groups subjected to stroke and treated after 3hr and for the following 4 days with daily IP injections of either vehicle or DMM solution (160 mg/Kg body weight). As illustrated in **Fig. 1**, the day of the stroke surgery was indicated as Day 0, and behavioural tests were performed 1 or 2 days before surgery (respectively −1 and −2) and at 48 h (day 2) and 96 h post-stroke (day 4, see **Supplementary Information**). On day 4, mice were sacrificed and their brains collected/stored overnight in ice-cold 10% formalin (v/v) before being washed with PBS and cut into transversal sections of 35 μm thickness using a vibratome (Leica VT 1200, see **Supplementary Information**).

## Acknowledgements

This research was supported by a Wellcome Trust Clinical Research Career Development Fellowship G105713 (LPJ) and the Intramural Research Program of the NINDS/NIH (JDB and JH). Work in the MPM lab is supported by the Medical Research Council UK (MC_UU_00028/4) and by a Wellcome Trust Investigator award (220257/Z/20/Z).

## Author contributions

JDB and LPJ conceptualized the project.

JDB designed the experimental protocols.

JDB and YJL performed the studies.

TL performed bioinformatics analysis.

MEGS performed the metabolomic analysis.

EG performed analysis of gene expression data and integration with metabolomics dataset.

CMW, AMN, and LWT performed the *in vitro* experiments and analysis.

DA, MRB, and MC performed *in vivo* experiments and analysis.

JDB, CMW, MEGS, NV, and LPJ analysed and interpreted all data.

JDB, AC, FG, SI, JS, SS, MPM, MA, MC, and LPJ wrote the manuscript with assistance/contributions from all the other authors.

JDB, MPM, MA, MC, JMH, and LPJ provided supervision/guidance to all parties.

## Competing interests

JDB and EG have an equity positions POCKiT Diagnostics which is developing a diagnostic tool for LVO detection. JDB and JS have an equity positions in Centile Bioscience. JDB also hold positions/equity in Treovir and NeuroX1. MPM holds patents in the therapeutic application of malonates.

## Extended Data Figures

**Extended Data Figure 1.**
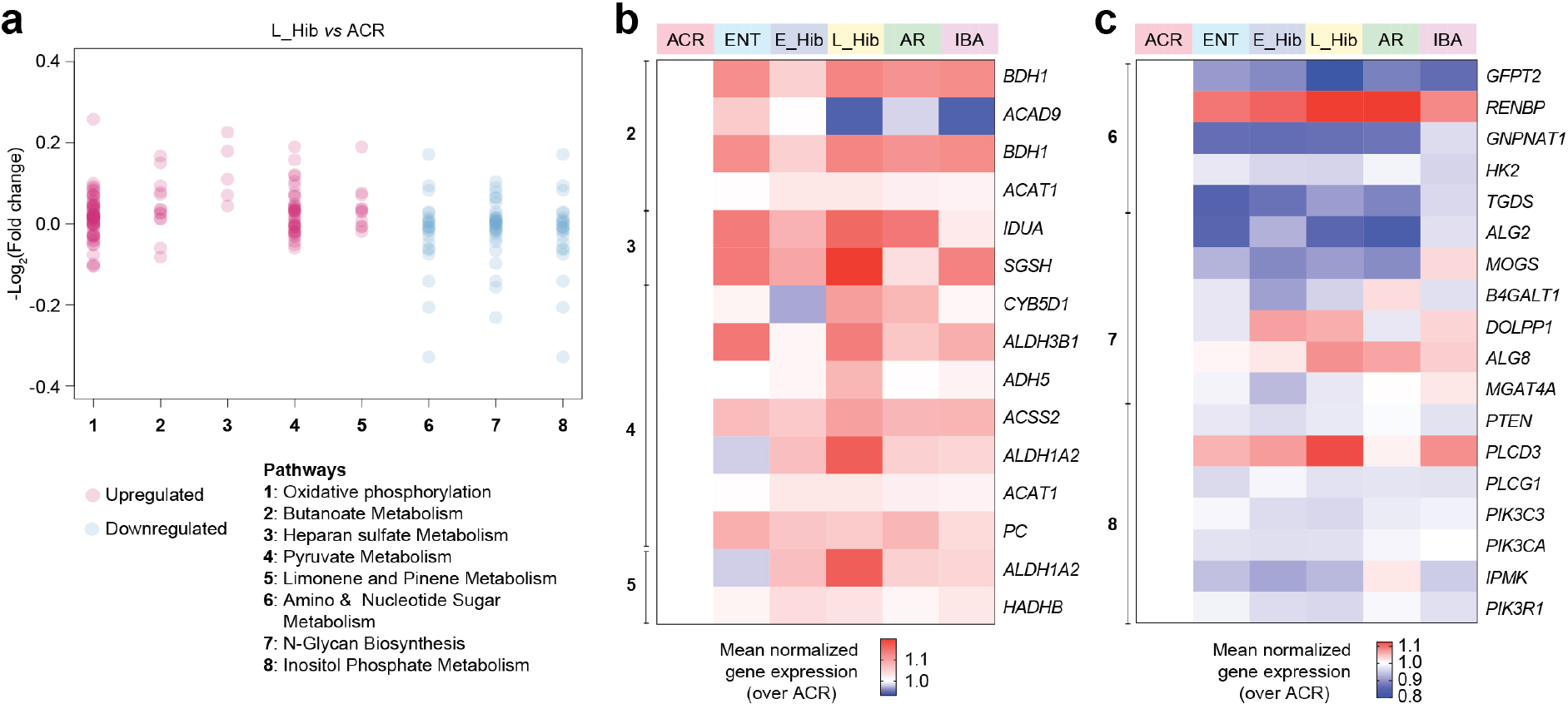
Genes determining the GSEA pathways of L_Hib *vs* ACR. **a**, Significantly enriched metabolic pathways according to GSEA analysis performed between L_Hib and ACR. Single dots are all single genes. **b**, Heat map of selected metabolic genes (P-value <0.1) defining the GSEA pathways upregulated in L_Hib *vs* ACR (except oxidative phosphorylation, reported in **Fig. 1**) at defined stages of hibernation. **c**, Heat map of selected metabolic genes (P-value < 0.1) defining the GSEA pathways downregulated in L_Hib *vs* ACR at defined stages of hibernation. Abbreviations: ACR (active cold room), ENT (entrance of hibernation), E_Hib (early hibernation), L_Hib (late hibernation/torpor), AR (arousal), IBA (interbout arousal).

**Extended Data Figure 2.**
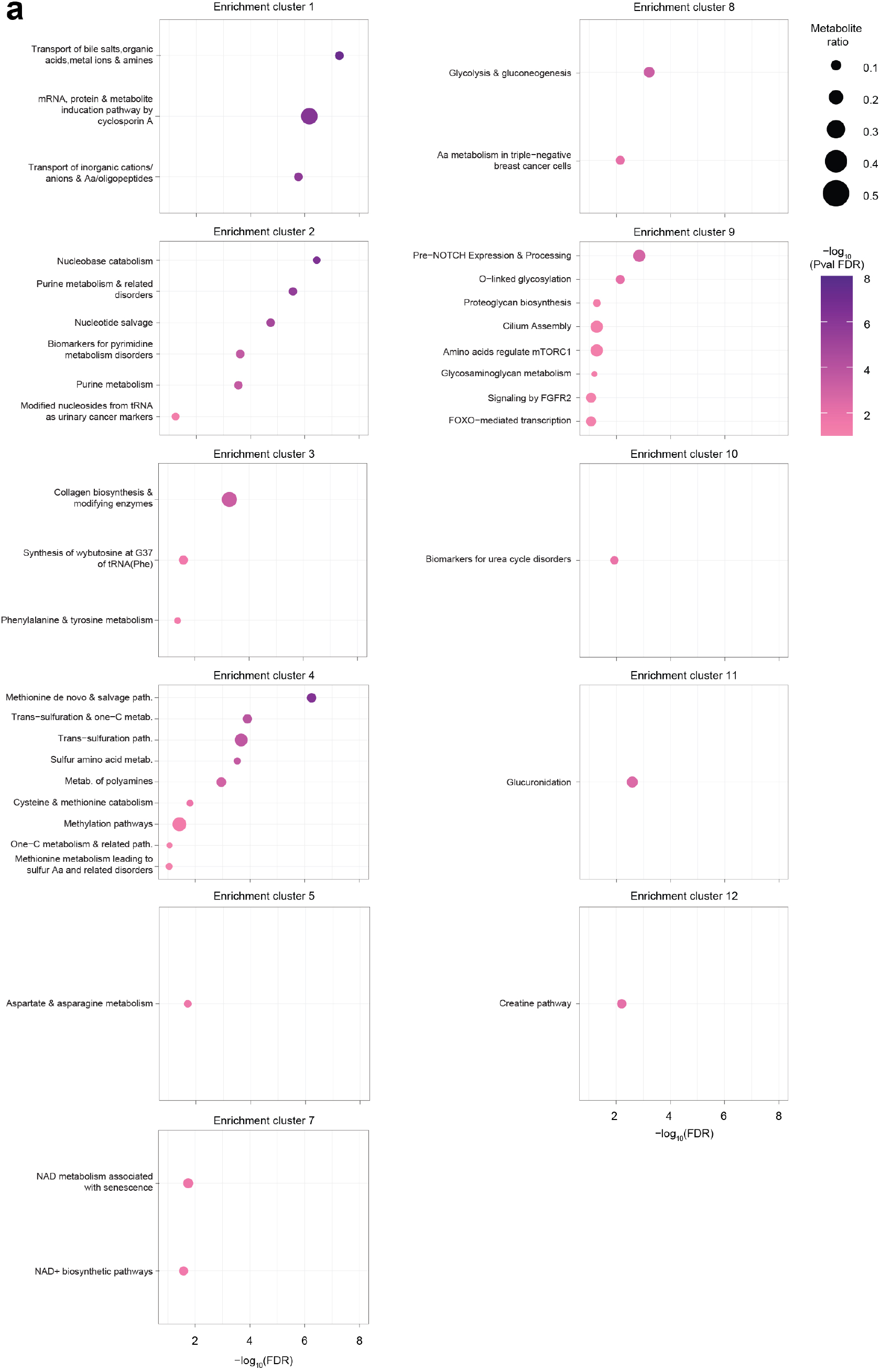
Custers derived from pathway enrichment analysis. **a,** Enrichment of WIKI pathways of the metabolic enrichment clusters (except *enrichment cluster 6*, reported in **Fig. 2**). Circle size is proportional to the metabolite ratio in the pathway and colorimetric scale shows the −log10 value of significance after Benjamini-Hochberg (BH) correction for multiple testing or false discovery rate. n=6 biological replicates per group.

**Extended Data Figure 3.**
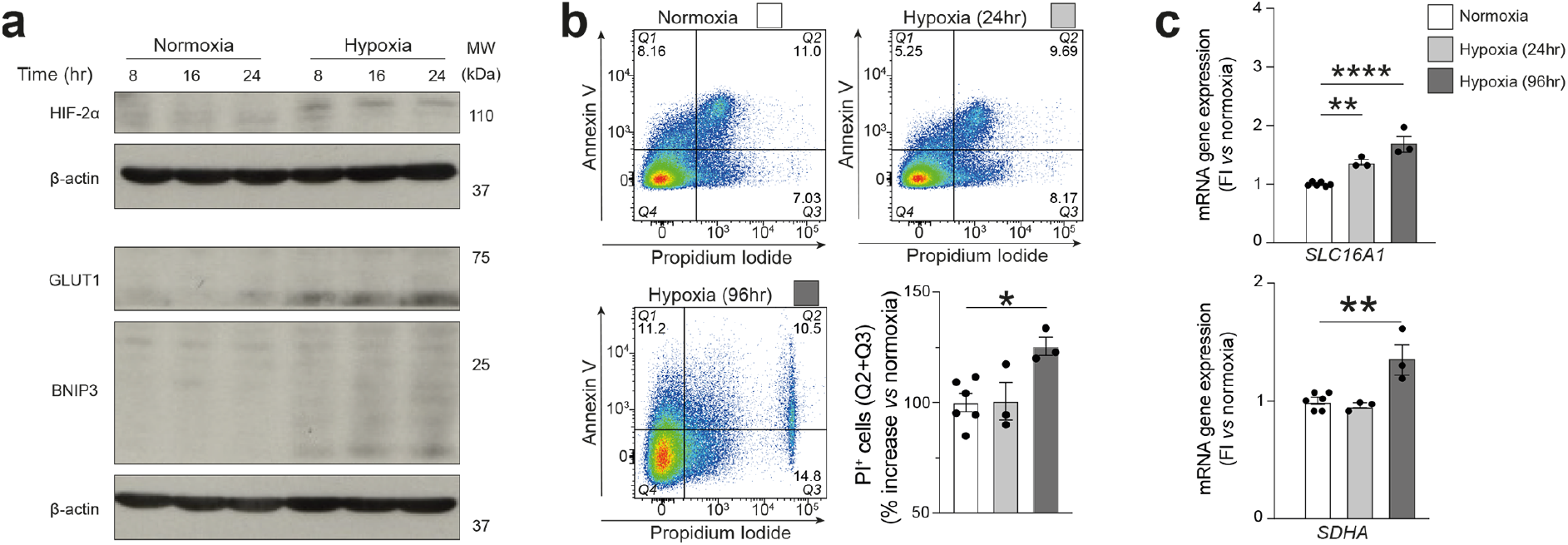
*In vitro* model of hypoxia in human SH-SY5Y neurons. **a**, Western blot showing the increased induction of the hypoxic markers HIF-2α, GLUT1, and BNIP3 at increasing times of hypoxia (1% O_2_) *vs* normoxia (21% O_2_). HIF-1α protein induction was also detectable (not shown). **b**, Annexin/PI staining of SH-SY5Y neurons at 24 and 96 h of hypoxia. n=3-6 technical replicates per group. Significance was determined via one-way ANOVA followed by Dunnet’s multiple comparisons test for correction. *: p<0.05. **c**, Gene expression analysis of selected hypoxic genes found to be increased during hibernation (i.e., *SLC16A1, SDHA*) in SH-SY5Y neurons at 24 and 96 h of hypoxia. n=3-6 technical replicates per group. Significance was determined via one-way ANOVA followed by Dunnet’s multiple comparisons test for correction. *: p<0.05; **: p<0.01, ****: p<0.0001.

## Supplementary Methods

### *Ex vivo* Ultrahigh Performance Liquid Chromatography-Tandem Mass Spectroscopy (UPLC-MS/MS)

Untargeted metabolomics profiling of brain samples was performed by Metabolon (Durham, NC)^1^.Samples were prepared using the automated MicroLab STAR^®^ system from Hamilton Company. Metabolite extraction, along with several recovery standards for QC purposes, was performed via methanol precipitation under vigorous shaking for 2 min (Glen Mills GenoGrinder 2000) followed by centrifugation. The resulting extract was divided into fractions for analysis by separate reverse phase (RP)/UPLC-MS/MS methods with positive ion mode electrospray ionization (ESI), RP/UPLC-MS/MS with negative ion mode ESI, HILIC/UPLC-MS/MS with negative ion mode ESI, and backup. A pooled matrix sample as well as extracted water samples served as technical replicate and process blanks throughout runs. A cocktail of known QC were also spiked into every analysed sample for instrument performance monitoring and chromatographic alignment. Experimental samples were randomized across the platform run with QC samples spaced evenly among the injections. All methods utilized a Waters ACQUITY ultra-performance liquid chromatography (UPLC) and a Thermo Scientific Q-Exactive high resolution/accurate mass spectrometer interfaced with a heated electrospray ionization (HESI-II) source and Orbitrap mass analyzer operated at 35,000 mass resolution. Compounds were identified by comparison to library entries of containing purified standards or additional compounds validated by Metabolon via data curation. Metabolites were identified based on their retention time/index (RI), mass to charge ratio (*m/z*), and chromatographic data (including MS/MS spectral data) as compared to all other molecules present in the Metabolon library. Sucesfull matches were determined based on the presence of a narrow RI window of the proposed identification, accurate mass match to the library ± 10 ppm, and the MS/MS forward and reverse scores between the experimental and authentic standards spectra for a given metabolite. Peaks were quantified using area-under-the-curve. Day-to-day variability was accounted by registering the metabolite medians to one and normalizing each metabolite within a particular sample proportionately to their median. The resulting raw intensities across all samples are found in Table 2a.

### Metabolomics processing and statistical analysis

Data processing of integrated peak intensities was performed in the R environment, version 3.6.2 (https://www.R-project.org/). Briefly, metabolites with missing values (N/As) in 33% or more of the samples were removed, with the remaining N/As imputed using the k-Nearest Neighbour algorithm as implemented on the package *VIM* (version 5.1.1)^2^. Data underwent log-10 transformation and autoscale normalization using core functions in the R environment; normalized intensities were graphically inspected to verify appropriate normalization. The resulting normalized intensities across all samples are found in Table 2b. All samples were included in the analyses and no samples or animals were excluded. Multivariate statistical analysis was initially performed on the normalized dataset using the Sparse Partial Least Squares – Discriminant Analysis (sPLS-DA) algorithm as implemented in the R package mixOmics (version 6.10.9)^3^. sPLS-DA models allow for the identification of variance attributed to class membership, while selecting variables meaningful to the classification task^4^. Metabolites with a variable influence of projection (VIP) > 1 in Component 1 were retained for further analyses due to their importance in contributing to the model’s classification task^5^. Hierarchical clustering and heatmaps of features were calculated and generated using the package *pheatmap* (version 1.0.12); metabolic pathway annotation within heatmap was performed using Metabolon’s custom annotation panel. Univariate analysis of features with VIP > 1 was performed using core functions in R; metabolites underwent Kruskal-Wallis nonparametric testing followed by false discovery rate (FDR) correction via the Benjamini-Hochberg (BH) method. Features were considered statistically significant if P-value < 0.05 upon FDR correction and quantitative data are presented box and whisker plots (min max).

### Metabolic pathway enrichment analysis

Pathway enrichment analysis of metabolites with a VIP > 1 in Component 1 was performed using the Relational Database of Metabolic Pathways^6^, using WikiPathways v20220710 as data source^7^. Significance was determined with right-tailed Fischer’s exact test followed by FDR correction via the BH method (α = 0.050). Metabolite ratio was defined across enriched pathways as the number of mapped input metabolites in relation to the total number of metabolites annotated in a given pathway^8^. Enriched pathways subsequently underwent Heuristic Multiple Linkage clustering based on partial overlap of the pathway’s mapped metabolites (Braisted et al 2023). Clusters metabolite ratio and FDR adjusted P-value were calculated using the median metabolite ratio and FDR adjusted P-value across pathways assigned to a given cluster.

### *Ex vivo* RNA sequencing

Fastq files were quality inspected by sample using the FASTQC tool (http://www.bioinformatics.babraham.ac.uk/projects/fastqc/). Adaptor clipping and quality trimming was achieved using Trimmomatic (http://www.usadellab.org/cms/index.php?page=trimmomatic). Stranded paired-end reference mapping against the current instance of the Squirrel genome (spetri2) via local alignment was performed in CLCbio (www.clcbio.com); providing RPKM expression for 20,387 transcripts/sample. Post enumeration, RPKM expression was imported into R (www.cran.r-project.org) and cross-sample normalized via quantile tact after pedestalling by 2 and taking the Log2 transform. Quality of the expression enumerated per sample was challenged and assured via Tukey box plot, covariance-based PCA scatter plot, and correlation-based heat map. To remove noise-biased expression, lowess fitting was employed to define the minimum threshold of expression at which the linear relationship between mean expression (i.e., signal) and CV (i.e., noise) across groups is grossly lost. Transcripts not having expression greater than this threshold for at least one sample were discarded with the remaining transcripts subject to ANOVA testing. Transcripts having an uncorrected P < 0.05 by this test were subset and post hoc analysis performed using the TukeyHSD test. Transcripts with a P < 0.05 by this test and an absolute difference of means >= 1.5X for a group comparison were subset as those having differential expression between those groups. Transcript annotations were obtained from the Ensembl FTP site (http://useast.ensembl.org/info/data/ftp/index.html). Transcript sequences were aligned against the *Ictidomys tridecemlineatus’s* genome. For the purpose of functional annotation, squirrel gene symbols from Ensembl (Table 1) were matched to the human counterpart to retrieve WikiPathways metabolic pathways annotations.

### Integration of RNA sequencing and metabolomics

All samples were included in the analyses and no samples or animals were excluded. Multivariate statistical analysis was initially performed on the RPKM values using the Sparse Partial Least Squares – Discriminant Analysis (sPLS-DA) algorithm as implemented in the R package mixOmics (version 6.10.9, Rohart F et al 2017). To define the biological pathways associated with each stage of hibernation, the log2 fold change between the median of each hibernation stage and the full dataset median expression was calculated. For each hibernation stage, this was used as input for a gene set enrichment analysis (GSEA) against the WikiPathways collection, which was downloaded from MsigDB using the R package ‘msigdbr’ version 7.5.1. For all GSEA analyses, the R package ‘piano’ version 2.14.0 was used. To focus on the metabolic gene expression we used a previously defined manually curated subset of metabolic genes and pathways^9^. To integrate the gene expression and metabolomics datasets, genes that significantly contributed to the pathway enrichment found by GSEA and metabolites belonging to *encrichment cluster 6* of the metabolomics sPLSDA analysis were selected and enzyme commission (EC) numbers associated to each gene and/or metabolite were extracted from the UniProt database. A hive plot was then built by linking each gene or metabolite to the associated EC number. The hive plot was built by using the R package ‘HiveR’ version 0.3.63.

### Succinate dehydrogenase (SDH) activity *in vitro*

To assess whether dimethyl malonate (DMM, Merck) could inhibit succinate dehydrogenase activity in SH-SY5Y cells, we used a fumarate detection kit (Abcam) following manufacturer’s protocol. Briefly, SH-SY5Y cells were seeded at a final density of 5×10^5^ cells/well in 6-well plates. After 9 days of differentiation, cells were treated with either 5 mM or 10 mM of DMM for 1 h in normoxic conditions at 37°C with 5% CO_2_. After 1 h, cells were washed with PBS and homogenized in the assay buffer provided. The assay was performed on 50 μl of sample per well in a new 96-well flat bottom culture plate. A standard curve (0-2-4-6-8-10 nmol/well) was obtained following developer’s recommendation and addding to each well a final volume adjusted to 50 μl, using assay buffer. The reaction mix was prepared and 100 μl was added to each well. The plate was incubated for 60 min in a 0% CO2 incubator at 37°C. After 60 min, the absorbance was measured at 450 nm with a Tecan Infinite M200 Pro plate reader. Data was analysed according to the manufacturer’s recommendations.

### SH-SY5Y *in vitro* culture

SH-SY5Y cells (gift from Michael Whitehead, University of Cambridge) were kept in culture and expanded in growth medium [DMEM-F12 (Gibco), 10% FBS, 1% pen/strep]. Growth medium was refreshed every 4 to 7 days. After reaching 80-90% confluency, SH-SY5Y cells were washed with PBS (Gibco), detached with trypsin-EDTA 0.05%, (Thermo Fisher Scientific) for 5 min followed by trypsin neutralization in growth media and spun down at 400 *g* for 5 min. Next, SH-SY5Y cells were counted and seeded at a density of 5×10^5^ cells/well for 6-well plates and 5×10^4^ cells/well on coverslips in 12-well plates with differentiation media (Neurobasal medium [Thermo Fisher Scientific], B27 supplement 2% [Thermo Fisher Scientific], GlutaMAX 1% [Thermo Fisher Scientific], and all-trans retinoic acid 10 *μ* M [STEMCELL Technologies]). Differentiation media was replaced every other day until day 9 post-seeding.

Hypoxia was achieved by incubating cells in 1% O_2_, 5% CO_2_, and 94% N_2_ in a Don Whitley H35 workstation. To assess cellular apoptosis under hypoxic conditions, we used an annexin V staining kit (eBioscience) in tandem with the viability dye propidium iodided (PI) followed by flow cytometry analysis. 5×10^5^ SH-SY5Y cells/well (n=3 technical replicates per condition) were differentiated for 9 days and then subjected to hypoxia (1% O_2_) for 24 h and 96 h. After 96 h, the conditioned media from each well was collected to capture all detached cells. Then, the remaining cells were detached using trypsin-EDTA 0.05% for 5 min and collected into the same tube as the conditioned media. The cells were spun down at 500 *g* for 5 min, the supernatant removed, and the pellet resuspended in binding buffer. The cells were then spun down at 500 *g* for 5 min, the supernatant removed, and the pellet resuspended in 100 μL of binding buffer plus 5 μL of annexin V antibody. The cells were incubated for 15 min at room temperature (RT) followed by pelleting at 500 *g* for 5 min, the supernatant removed, and washed in binding buffer followed by a final spin at 500 *g* for 5 min. The cell pellet was resuspended in 200 μL of binding buffer and stored on ice until acquisition. Acquisition was performed on a BD LSRFortessa™ flow cytometer (BD Biosciences) using the 640 nm red laser for annexin V and the 561 nm yellow laser for PI. Positive gates were established using unstained, live/dead, and single marker cells prior to the analysis of the samples. Immediately prior to acquisition, 5 μL of PI was added to each sample. 100,000 total events were acquired for each sample and analysis was performed in FlowJo v10 software (BD Biosciences).

For gene expression analysis, 5×10^5^ SH-SY5Y cells/well (n=3 technical replicates / condition) were differentiated for 9 days and then subjected to hypoxia (1% O_2_) for 24 h and 96 h. Total RNA from SH-SY5Y cells was collected by washing the cells once in ice-cold PBS followed by the addition of 350 μL of RLT lysis buffer (QIAGEN). Samples were then stored at −80°C until extraction. Total RNA from SH-SY5Y was extracted using the RNeasy Mini Kit (QIAGEN) following the manufacturer’s instructions. Total RNA was then quantified with the NanoDrop 2000c instrument (Thermo Fisher Scientific). For qRT-PCR analysis, 500 ng of RNA were reversed-transcribed using the high-capacity cDNA reverse transcription kit (Thermo Fisher Scientific) according to the manufacturer’s instructions using a 20 μL reaction volume. qRT-PCR was performed with the TaqMan™ Fast Universal PCR Master Mix (2x) (Thermo Fisher Scientific) and TaqMan^®^ Gene Expression Assays for: *HCAR1* (Thermo Fisher Scientific, Hs02597779_s1), *LDHB* (Thermo Fisher Scientific, Hs00929956_m1), *SLC16A1* (Thermo Fisher Scientific, Hs01560299_m1), *SDHA* (Thermo Fisher Scientific, Hs00188166_m1), *NAMPT* (Thermo Fisher Scientific, Hs00237184_m1), and 18S (Thermo Fisher Scientific) was used for normalization. For qRT-PCR, 1.5 μL of cDNA from each sample was run in triplicates using a 7500 Fast Real-Time PCR System (Applied Biosystems) and analysed with the 2^-ΔΔCT^ method.

For neurite length, 5×10^4^ SH-SY5Y cells/well were plated onto 12 mm glass coverslips in 12-well plates and differentiated for 9 days. The cells were then subjected to hypoxia (1% O2) in the presence or absence of 10 mM DMM for 24 h. Immediately after, the coverslips were washed once in PBS followed by fixation in 4% PFA for 60 mins at RT. The extended fixation was to ensure preservation of neurites for quantification. After fixation, the coverslips were washed three times in PBS, permeablized in PBS with 0.1% Triton™ X-100 (Merck) for 15 min, blocked with PBS with 5% normal goat serum for 30 min, then stained with phalloidin (Thermo Fisher Scientific) diluted 1:400 in PBS for 30 min followed by counterstaining with DAPI (Merck) for 10 min to label nuclei. The cells were then washed three times in PBS and mounted onto glass slides using ProLong™ diamond antifade mountant (Thermo Fisher Scientific). Slides were left to dry overnight and stored at 4°C until acquisition. Bright field pictures (10x) of 5 ROIs per coverslip from each condition were taken using a Leica DM IL LED microscope. Images were converted to 8-bit and contrast was adjusted so the neurites were easily visible using Fiji 2.0.0. software. Data were quantified using the NeuronJ plugin that allows for semi-automatic tracing from an average of >30 neurite lengths per ROI from n=3 independent experiments.

### Western blotting

Protein lysates were generated by rapid (>2 min) lysis of cells using Laemmli sample buffer (2% SDS, 10% glycerol, 10% 0.625 M Tris-HCl pH 6.8, in water), followed by heating to 100 °C for 5 min. Protein concentration in lysates was determined by BCA assay (ThermoFisher Scientific, #23227), and lysates were normalised to equal protein concentration by addition of Laemmli sample buffer. Lysates were prepared for western blotting by the addition of 0.4% (w/v) bromophenol blue solution (10% final v/v), and 1M dithiothreitol (10% final v/v).

Samples were separated by gel electrophoresis using a Hoefer SE400 vertical protein electrophoresis unit, then transferred to PVDF membranes using a Hoefer TE42 transfer unit. Membranes were blocked with 5 % (w/v) non-fat dry milk in TBS-Tween (20 mM Tris, 150 mM NaCl, 0.1% (v/v) Tween-20), for 1 h at RT. Proteins were detected by serial incubation with target-specific antibodies (16-24 h, 4 °C), followed by species-specific HRP-linked secondary antibodies (1 h, RT), then incubation with enhanced chemiluminescence substrate (Cytiva, #RPN2209). HIF-1α and HIF-2α were exposed for 30 min on x-ray film to obtain protein bands with GLUT1 and BNIP3 exposed for 5 min. Antibody details as follow: mouse anti-human HIF-1α (BD Biosciences, #610959); rabbit anti-human HIF-2α (Cell Signaling Technologies, #7096); rabbit anti-human GLUT1 (Cell Signaling Technologies, #73015); rabbit anti-human BNIP3 (Cambridge Biosciences, #HPA003015); mouse anti-human β-actin (Abcam, #ab6276).

### Photothrombotic stroke induction

Before surgery, male C57 mice (8-10 weeks of age) were anesthetized via intraperitoneal injection of 80 mg/kg ketamine and 5mg/kg xylazine. Anesthetized mice were placed in a stereotaxic frame and their body temperature was maintained at 37 °C by a digital heating mat. After removing the skin, skull was exposed and the green laser (λ 530 nm) was positioned in the M1 forelimb area (from bregma: 0,4 mm AP and ± 1,6 mm ML) contralateral to the preferred paw (see also Behavioural tests section below). 200 μl of Rose Bengal solution (10 mg/ml in physiological solution) was delivered via intraperitoneal injection and after 5 min the laser was switched on (33 mW) and M1 irradiated for 15 min.

### Nissl staining, images acquisition and ischemic volume measurement

For ischemic volume measurements, one of every three brain sections of 35 μm each was stained with 0.1 % cresyl violet (Nissl staining). Briefly, slides were submerged for 15 min in a mixture of absolute ethanol and chloroform (in 4:1 ratio), hydrated for 2 min in MilliQ water, stained for 2 min in 0.1% cresyl violet solution and dehydrated by quick passages through 70%, 80% and 96% ethanol solutions. Finally, slides were cleared in absolute xylene for 10 min following a washing step in 96% ethanol before being mounted using Eukitt mounting medium (#0389, Merck).

Images were acquired on a Leica bright bright-field microscope (Leica DM6000) using a 5X objective and ischemic volume quantified in Fiji software as follows: ischemic volume (mm^3^) = area (mm^2^) * section thickness * spacing factor

### Behavioural tests

To assess wether DMM may influence motor recovery after stroke, we performed a variety of behavioural tests as has been previously decribed^10^:

#### Cylinder test

The cylinder test was used to evaluate the spontaneous asymmetry in forelimb use during vertical-lateral wall exploration of a clear plexiglass cylinder (10 cm diameter and 14 cm in height). Mice were individually recorded for 5 min with a video camera placed below the cylinder and the preferential use of the right or left forelimb was scored by counting the number of times an individual forepaw was used by mice to rear their body from the bottom of the cylinder, to touch the cylinder wall, to push themselves from a region to another of the wall as well as down from the wall, and to land on the bottom of the cylinder. Simultaneous use of both left and right forelimb was not taken into consideration in the counting. We evaluated the forelimb preference first at baseline (i.e. prior to photothrombotic stroke) in the forelimb motor cortex contralateral to the preferred paw, and next at 48 h (2 days) and 96 h (4 days) after stroke induction. The asymmetry index was subsequently calculated trough the following formula: (C_ipsi_/C_ipsi_ + C_contra_) × 100 − (C_contra_/C_ipsi_ + C_contra_) × 100, where the Cipsi represents the ipsilateral paw to the hemisphere subjected to injury.

#### Gridwalk test

The gridwalk test was performed to assess motor fine function and coordination in mice following stroke and DMM treatments. Mice were placed on a metal grid (25 cm length and 25 cm width) composed of wire-openings (1 × 1) to explore freely for 5 min. The number of correct steps and foot faults were video-recorded by a camera placed underneath the grid, and steps made using the preferential forelimb were counted via analyzing the video in VLC Media Player. We considered a step failure when the the foot slipped into a grid hole and percentage of foot faults was calculated according to the formula: Foot Faults% = 100 × #foot faults/#correct steps + #foot fault. Values obtained at 48 h and 96 h were normalized to baseline. Rotarod test. To assess motor coordination and function, mice were placed on a horizontal rotating rod with 3.5 cm diameter using the accelerating rotation speed mode (4–40 rpm within 5 min). The trial was performed twice with an inter-trial interval of 20 min and stopped when the mice fell down. The latency to fall (in terms of seconds) was recorded and the best performance for each mouse was used for statistical analysis. Trials were performed at baseline and at 48 h and 96 h after stroke induction, and values normalized to baseline. For better performance, mice were pre-trained on the rotarod before baseline at a fixed speed of 8 rpm until the score was at least 60 seconds.

Open field. The open field test was carried out to assess spontaneous and and exploratory activity in mice after stroke and following DMM treatments. Each mouse was gently placed in the center a square wooden box (47 cm × 47 cm) and its explorative activity was recorded under low light conditions for 5 min using a camera. Total distance traveled, time spent in center and number center entries were analyzed using VideoTrack software. All values obtained at 48 h and 96 h were normalized to baseline.

### Other statistical analyses

Statistically significant differences between groups were analysed using a One-Way ANOVA for multiple comparisons, or a two-way unpaired Student’s t-test for comparisons of two groups. Differences were considered significant at P-value < 0.05 and quantitative data are presented as box and whisker plots (min/max).

## References

1 Deb, P., Sharma, S. & Hassan, K. M. Pathophysiologic mechanisms of acute ischemic stroke: An overview with emphasis on therapeutic significance beyond thrombolysis. Pathophysiology: the official journal of the International Society for Pathophysiology / ISP 17, 197–218, doi:10.1016/j.pathophys.2009.12.001 (2010).

2 Frerichs, K. U., Kennedy, C., Sokoloff, L. & Hallenbeck, J. M. Local cerebral blood flow during hibernation, a model of natural tolerance to “cerebral ischemia”. J Cereb Blood Flow Metab 14, 193–205, doi:10.1038/jcbfm.1994.26 (1994).

3 Benjamin, E. J. et al. Heart Disease and Stroke Statistics-2019 Update: A Report From the American Heart Association. Circulation 139, e56–e528, doi:10.1161/CIR.0000000000000659 (2019).

4 Lo, E. H., Moskowitz, M. A. & Jacobs, T. P. Exciting, radical, suicidal: how brain cells die after stroke. Stroke 36, 189–192, doi:10.1161/01.STR.0000153069.96296.fd (2005).

5 Wang, L. C. et al. The “hibernation induction trigger”: specificity and validity of bioassay using the 13-lined ground squirrel. Cryobiology 25, 355–362, doi:10.1016/0011-2240(88)90043-0 (1988).

6 Ginsberg, M. D. Adventures in the pathophysiology of brain ischemia: penumbra, gene expression, neuroprotection: the 2002 Thomas Willis Lecture. Stroke 34, 214–223, doi:10.1161/01.str.0000048846.09677.62 (2003).

7 Ma, Y. L. et al. Absence of cellular stress in brain after hypoxia induced by arousal from hibernation in Arctic ground squirrels. American journal of physiology. Regulatory, integrative and comparative physiology 289, R1297–1306, doi:10.1152/ajpregu.00260.2005 (2005).

8 Zhou, F. et al. Hibernation, a model of neuroprotection. Am J Pathol 158, 2145–2151, doi:10.1016/S0002-9440(10)64686-X (2001).

9 Na, U. et al. The LYR factors SDHAF1 and SDHAF3 mediate maturation of the iron-sulfur subunit of succinate dehydrogenase. Cell Metab 20, 253–266, doi:10.1016/j.cmet.2014.05.014 (2014).

10 Karp, P. D., Midford, P. E., Caspi, R. & Khodursky, A. Pathway size matters: the influence of pathway granularity on over-representation (enrichment analysis) statistics. BMC Genomics 22, 191, doi:10.1186/s12864-021-07502-8 (2021).

11 Toyoshima, S., Watanabe, F., Saido, H., Miyatake, K. & Nakano, Y. Methylmalonic acid inhibits respiration in rat liver mitochondria. J Nutr 125, 2846–2850, doi:10.1093/jn/125.11.2846 (1995).

12 Dutra, J. C. et al. Inhibition of succinate dehydrogenase and beta-hydroxybutyrate dehydrogenase activities by methylmalonate in brain and liver of developing rats. J Inherit Metab Dis 16, 147–153, doi:10.1007/BF00711328 (1993).

13 Cordes, T. & Metallo, C. M. Itaconate Alters Succinate and Coenzyme A Metabolism via Inhibition of Mitochondrial Complex II and Methylmalonyl-CoA Mutase. Metabolites 11, doi:10.3390/metabo11020117 (2021).

14 Martinez-Reyes, I. & Chandel, N. S. Mitochondrial TCA cycle metabolites control physiology and disease. Nature communications 11, 102, doi:10.1038/s41467-019-13668-3 (2020).

15 Xu, J. et al. Inhibiting Succinate Dehydrogenase by Dimethyl Malonate Alleviates Brain Damage in a Rat Model of Cardiac Arrest. Neuroscience 393, 24–32, doi:10.1016/j.neuroscience.2018.09.041 (2018).

16 Uzdensky, A. B. Photothrombotic Stroke as a Model of Ischemic Stroke. Translational stroke research 9, 437–451, doi:10.1007/s12975-017-0593-8 (2018).

17 Cherchi, L., Anni, D., Buffelli, M. & Cambiaghi, M. Early Application of Ipsilateral Cathodal-tDCS in a Mouse Model of Brain Ischemia Results in Functional Improvement and Perilesional Microglia Modulation. Biomolecules 12, doi:10.3390/biom12040588 (2022).

18 Dark, J. Annual lipid cycles in hibernators: integration of physiology and behavior. Annu Rev Nutr 25, 469–497, doi:10.1146/annurev.nutr.25.050304.092514 (2005).

19 Frerichs, K. U. et al. Suppression of protein synthesis in brain during hibernation involves inhibition of protein initiation and elongation. Proc Natl Acad Sci U S A 95, 14511–14516, doi: 10.1073/pnas.95.24.14511 (1998).

20 Galster, W. & Morrison, P. R. Gluconeogenesis in arctic ground squirrels between periods of hibernation. Am J Physiol 228, 325–330, doi:10.1152/ajplegacy.1975.228.1.325 (1975).

21 D’Alessandro, A., Nemkov, T., Bogren, L. K., Martin, S. L. & Hansen, K. C. Comfortably Numb and Back: Plasma Metabolomics Reveals Biochemical Adaptations in the Hibernating 13-Lined Ground Squirrel. J Proteome Res 16, 958–969, doi:10.1021/acs.jproteome.6b00884 (2017).

22 Nelson, C. J., Otis, J. P., Martin, S. L. & Carey, H. V. Analysis of the hibernation cycle using LC-MS-based metabolomics in ground squirrel liver. Physiol Genomics 37, 43–51, doi:10.1152/physiolgenomics.90323.2008 (2009).

23 Serkova, N. J., Rose, J. C., Epperson, L. E., Carey, H. V. & Martin, S. L. Quantitative analysis of liver metabolites in three stages of the circannual hibernation cycle in 13-lined ground squirrels by NMR. Physiol Genomics 31, 15–24, doi:10.1152/physiolgenomics.00028.2007 (2007).

24 Rice, S. A. et al. Nitrogen recycling buffers against ammonia toxicity from skeletal muscle breakdown in hibernating arctic ground squirrels. Nat Metab 2, 1459–1471, doi:10.1038/s42255-020-00312-4 (2020).

25 Regan, M. D. et al. Nitrogen recycling via gut symbionts increases in ground squirrels over the hibernation season. Science 375, 460–463, doi:10.1126/science.abh2950 (2022).

26 Henry, P. G. et al. Brain energy metabolism and neurotransmission at near-freezing temperatures: in vivo (1)H MRS study of a hibernating mammal. J Neurochem 101, 1505–1515, doi:10.1111/j.1471-4159.2007.04514.x (2007).

27 Gonzalez-Riano, C. et al. Metabolomic Study of Hibernating Syrian Hamster Brains: In Search of Neuroprotective Agents. J Proteome Res 18, 1175–1190, doi:10.1021/acs.jproteome.8b00816 (2019).

28 Luan, Y. et al. Integrated transcriptomic and metabolomic analysis reveals adaptive changes of hibernating retinas. J Cell Physiol 233, 1434–1445, doi:10.1002/jcp.26030 (2018).

29 Hallenbeck, J. M. & Dutka, A. J. Background review and current concepts of reperfusion injury. Arch Neurol 47, 1245–1254, doi:10.1001/archneur.1990.00530110107027 (1990).

30 Drew, K. L. et al. Role of the antioxidant ascorbate in hibernation and warming from hibernation. Comp Biochem Physiol C Toxicol Pharmacol 133, 483–492, doi:10.1016/s1532-0456(02)00118-7 (2002).

31 Zhang, Z. et al. Protective effects of Dimethyl malonate on neuroinflammation and blood-brain barrier after ischemic stroke. Neuroreport 32, 1161–1169, doi:10.1097/WNR.0000000000001704 (2021).

32 Mottahedin, A. et al. Targeting succinate metabolism to decrease brain injury upon mechanical thrombectomy treatment of ischemic stroke. Redox Biol 59, 102600, doi:10.1016/j.redox.2023.102600 (2023).

33 Lee, Y. J. et al. Protein SUMOylation is massively increased in hibernation torpor and is critical for the cytoprotection provided by ischemic preconditioning and hypothermia in SHSY5Y cells. J Cereb Blood Flow Metab 27, 950–962, doi:10.1038/sj.jcbfm.9600395 (2007).

34 Lee, Y. J., Bernstock, J. D., Klimanis, D. & Hallenbeck, J. M. Akt Protein Kinase, miR-200/miR-182 Expression and Epithelial-Mesenchymal Transition Proteins in Hibernating Ground Squirrels. Front Mol Neurosci 11, 22, doi:10.3389/fnmol.2018.00022 (2018).

## References

1 Dehaven, C. D., Evans, A. M., Dai, H. & Lawton, K. A. Organization of GC/MS and LC/MS metabolomics data into chemical libraries. J Cheminform 2, 9, doi:10.1186/1758-2946-2-9 (2010).

2 Kowarik, A. & Templ, M. Imputation with the R Package VIM. Journal of Statistical Software 74, doi:10.18637/jss.v074.i07 (2016).

3 Rohart, F., Gautier, B., Singh, A. & Le Cao, K. A. mixOmics: An R package for ‘omics feature selection and multiple data integration. Plos Computational Biology 13, doi:ARTN e1005752 10.1371/journal.pcbi.1005752 (2017).

4 Cao, K. A. L., Boitard, S. & Besse, P. Sparse PLS discriminant analysis: biologically relevant feature selection and graphical displays for multiclass problems. Bmc Bioinformatics 12, doi:Artn 253 10.1186/1471-2105-12-253 (2011).

5 Liu, K. D. et al. Consequences of Lipid Remodeling of Adipocyte Membranes Being Functionally Distinct from Lipid Storage in Obesity. Journal of Proteome Research 19, 3919–3935, doi:10.1021/acs.jproteome.9b00894 (2020).

6 Braisted, J. et al. RaMP-DB 2.0: a renovated knowledgebase for deriving biological and chemical insight from metabolites, proteins, and genes. Bioinformatics, doi:ARTN btac726 10.1093/bioinformatics/btac726 (2022).

7 Slenter, D. N. et al. WikiPathways: a multifaceted pathway database bridging metabolomics to other omics research. Nucleic Acids Research 46, D661–D667, doi:10.1093/nar/gkx1064 (2018).

8 Garcia-Segura, M. E. et al. Pathway-based integration of multi-omics data reveals lipidomics alterations validated in an Alzheimer’s disease mouse model and risk loci carriers. Journal of Neurochemistry 164, 57–76, doi:10.1111/jnc.15719 (2023).

9 Gaude, E. & Frezza, C. Tissue-specific and convergent metabolic transformation of cancer correlates with metastatic potential and patient survival. Nature communications 7, 13041, doi:10.1038/ncomms13041 (2016).

10 Cherchi, L., Anni, D., Buffelli, M. & Cambiaghi, M. Early Application of Ipsilateral Cathodal-tDCS in a Mouse Model of Brain Ischemia Results in Functional Improvement and Perilesional Microglia Modulation. Biomolecules 12, doi:10.3390/biom12040588 (2022).

